# *Phytophthora infestans* RXLR-WY effector AVR3a associates with a Dynamin-Related Protein involved in endocytosis of a plant pattern recognition receptor

**DOI:** 10.1101/012963

**Authors:** Angela Chaparro-Garcia, Simon Schwizer, Jan Sklenar, Kentaro Yoshida, Jorunn I. B. Bos, Sebastian Schornack, Alexandra M. E. Jones, Tolga O. Bozkurt, Sophien Kamoun

## Abstract

Perception of pathogen associated molecular patterns (PAMPs) by cell surface localized pattern recognition receptors (PPRs), activates plant basal defense responses in a process known as PAMP/PRR-triggered immunity (PTI). In turn, pathogens deploy effector proteins that interfere with different steps in PTI signaling. However, our knowledge of PTI suppression by filamentous plant pathogens, i.e. fungi and oomycetes, remains fragmentary. Previous work revealed that BAK1/SERK3, a regulatory receptor of several PRRs, contributes to basal immunity against the Irish potato famine pathogen *Phytophthora infestans*. Moreover BAK1/SERK3 is required for the cell death induced by *P. infestans* elicitin INF1, a protein with characteristics of PAMPs. The *P. infestans* host-translocated RXLR-WY effector AVR3a is known to supress INF1-mediated defense by binding the E3 ligase CMPG1. In contrast, AVR3a^Kl-Y147del^, a deletion mutant of the C-terminal tyrosine of AVR3a, fails to bind CMPG1 and suppress INF1 cell death. Here we studied the extent to which AVR3a and its variants perturb additional BAK1/SERK3 dependent PTI responses using the plant PRR FLAGELLIN SENSING 2 (FLS2). We found that all tested variants of AVR3a, including AVR3a^Kl-Y147del^, suppress early defense responses triggered by the bacterial flagellin-derived peptide flg22 and reduce internalization of activated FLS2 from the plasma membrane without disturbing its nonactivated localization. Consistent with this effect of AVR3a on FLS2 endocytosis, we discovered that AVR3a associates with the Dynamin-Related Protein DRP2, a plant GTPase implicated in receptor-mediated endocytosis. Interestingly, DRP2 is required for ligand-induced FLS2 internalization but does not affect internalization of the growth receptor BRASSINOSTEROID INSENSITIVE 1 (BRI1). Furthermore, overexpression of DRP2 suppressed accumulation of reactive oxygen species triggered by PAMP treatment. We conclude that AVR3a associates with a key cellular trafficking and membrane-remodeling complex involved in immune receptor-mediated endocytosis and signaling. AVR3a is a multifunctional effector that can suppress BAK1/SERK3 mediated immunity through at least two different pathways.

**AUTHOR SUMMARY:** Plants have a basal layer of immunity to mount defense responses against invading pathogens; in turn, pathogens deploy effector proteins to subvert plant immunity and manipulate host processes to enable parasitic infection. The Irish potato famine pathogen *Phytophthora infestans* has a large set of effectors that target multiple host cellular sites. The best-characterized *P. infestans* effector AVR3a supports enhanced infection and suppresses the cell death induced by the *P. infestans* protein INF1-elicitin. Previous work demonstrated that the plant immunity co-receptor BAK1/SERK3 contributes to basal immunity to *P. infestans* and that the RXLR-WY effector of *P. infestans* suppresses BAK1/SERK3-mediated immunity by binding the E3 ligase protein CMPG1. Here we show that AVR3a suppresses additional defense responses mediated by BAK1/SERK3 independently of CMPG1. AVR3a reduces the endocytosis of the plant receptor FLS2, which recognizes the flagellin-derived peptide flg22 in a BAK1/SERK3 dependent manner. Furthermore, we demonstrate that AVR3a associates with the Dynamin-Related Protein DRP2, a plant GTPase involved in receptor-mediated endocytosis that is required for FLS2 internalization. Our work revealed that AVR3a is a multifunctional effector that perturbs cellular trafficking initiated at the cell periphery by at least two mechanisms, and that this effector associates with a key cellular trafficking and membrane-remodeling complex involved in immune receptor-mediated endocytosis and signaling.

## INTRODUCTION

Plants must respond in a timely and effective manner to external cues such as biotic and abiotic stimuli. In particular, plants are associated with a huge variety of microorganisms, many of which are sophisticated parasites^1–3^. To keep such parasitic microbes at bay, plants have evolved a multilayered immune system, which is triggered by microbial perception via receptors located at the cell surface and the cytoplasm^3, 4^. An important layer of defense relies on perception of pathogen associated molecular patterns (PAMPs) by cell surface localized pattern recognition receptors (PRRs), a basal defense response known as PAMP/PRR–triggered immunity (PTI)^5^. Early PTI signaling events include ion fluxes, reactive oxygen species (ROS) production, induction of mitogen-activated protein kinases (MAPKs) and calcium-dependent protein kinases (CDPKs) and transcriptional reprogramming^6–8^. Collectively, PTI contributes to immunity by delaying or arresting pathogen invasion, and in many cases PTI-deficient plant mutants became more susceptible to pathogens^9^. However, a common feature of adapted plant pathogens is the deployment of effector proteins that interfere with different steps in PTI signaling pathways^3, 4, 10, 11^. This is particularly striking for bacterial plant pathogens, which evolved a battery of often redundant effectors to counteract plant immunity elicited by conserved patterns such as flagellin and the elongation factor thermo unstable (EF-Tu)^3, 12, 13^. In contrast, our understanding of how eukaryotic pathogens, such as oomycetes and fungi, suppress PTI remains fragmentary^14–17^

The molecular signaling mechanisms activated after PAMP perception have been described for few PRRs. In *Arabidopsis thaliana* the leucine-rich repeat receptor-like kinases (LRR-RLKs) FLAGELLIN-SENSING 2 (FLS2) and EF-Tu RECEPTOR (EFR) recognize peptides derived from bacterial flagellin and elongation factor-Tu, respectively^18^. In addition, the CHITIN ELICITOR RECEPTOR KINASE 1 (CERK1) and the LYSIN-MOTIF RECEPTOR-LIKE KINASE 5 (LYK5), together mediate binding and recognition of fungal chitin^19–24^. Activation of EFR and FLS2 leads to the recruitment of the regulatory LRR-RLK BRASSINOSTEROID INSENSITIVE 1 (BRI1)-associated receptor kinase (BAK1, also called SERK3) to activate signal transduction^25–29^. Interestingly, BAK1/SERK3 is a positive regulator and interacts with the RLK BRI1 (BRASSINOSTEROID INSENSITIVE 1) to regulate brassinosteroid signaling and plant growth^30–32^, suggesting that common regulatory mechanisms between plant growth and immunity^33^.

BAK1/SERK3 is also required for defense responses mediated by the receptor like proteins (RLPs) Ve1, which confers resistance to race 1 strains of the fungal pathogens *Verticillium dahliae* and *Verticillium albo-atrum*, and LeEix2, which recognizes the fungal protein elicitor EIX (ETHYLENE-INDUCING XYLANASE)^34, 35^. More recently, BAK1/SERK3 emerged as a key component of basal defense against oomycetes, an important group of filamentous eukaryotic pathogens. Silencing of BAK1/SERK3 resulted in dramatically enhanced susceptibility and faster host colonization by the potato late blight pathogen *Phytophthora infestans*^36^. In addition, BAK1/SERK3 and its closest homolog BKK1/SERK4 are involved in resistance to the obligate biotrophic oomycete *Hyaloperonospora arabidopsidis*^28^ providing further evidence of the role of BAK1/SERK3-dependent basal resistance against oomycete pathogens. BAK1/SERK3 contribution to basal defense against oomycetes is most likely the result of recognition of PAMPs^26, 36, 37^. Indeed, BAK1/SERK3 is required for the cell death response triggered by INF1, a secreted *P. infestans* protein with features of PAMPs^26, 36^.

Plant endocytic trafficking has emerged as a dynamic process that is diverted towards sites of pathogen infection^38, 39^ indicating that this plant process may play a critical role in immune responses^40^. To maintain its protein levels at the cell surface, FLS2 undergoes constitutive endocytosis and recycles between plasma membrane and trans-Golgi network, independently of BAK1^25, 41^. However, FLS2 undergoes ligand-induced translocation into endosomal vesicles in a BAK1-dependent manner^41–44^. Therefore, FLS2 traffics through two different endocytic pathways depending on its activation status. Furthermore, BRI1 and BAK1/SERK3 also undergo constitutive recycling via endosomes and can localize to overlapping endosomal compartments^32^. Receptor-mediated endocytosis was initially thought as a mechanism for attenuation of signaling through depletion of activated receptor complexes. Further studies in animal systems revealed that receptor internalization contributes to additional signaling at the endocytic compartments^45^. In animal cells, late endosomal compartments, which host internalized receptors, regulate signaling events such as pro-inflammatory signaling, growth, and development^45–49^. In plants, localization of BRI1 and BAK1/SERK3 as well as a brassinosteroid analog at the same endosomal compartments pointed to a possible link between internalization and signaling in growth regulation^32, 50, 51^. However, defense-related endocytic signaling has yet to be unambiguously demonstrated in plants.

Mechanisms of receptor-mediated endocytosis involve clathrin-mediated and clathrin-independent pathways resulting in the recruitment of plasma membrane cargo^52–54^ that is later invaginated and pinched off into the cytoplasm often by the action of the large GTPase dynamin^53, 55^. Dynamin is a ~100 kDa protein that self-assembles into rings and helices to promote structural reorganization to mediate membrane fission in a reaction that requires GTP hydrolysis^55^. The mechanistic details of clathrin-mediated endocytosis have been well established in animal cells and dynamin has been implicated in the internalization of the immunity-related Interleukin-2 Receptor (IL-2R)^56^. However, in plants, mechanisms of endocytosis have not been explicitly studied^57, 58^ and the extent to which dynamin or dynamin-like proteins play a role in receptor mediated endocytosis or plant immunity remains poorly known.

*P. infestans* is the causal agent of potato and tomato late blight and a major threat to food security^59^. This oomycete pathogen deploys a large set of effectors that target multiple host cellular sites. Cytoplasmic effectors include the RXLR class, modular proteins that translocate inside host cells through poorly known mechanisms^60^. The biochemical activity of RXLR effectors is carried out by their C-terminal domains, which often contain variations of the conserved WY-domain fold^61, 62^. One example of RXLR-WY effector is AVR3a of *P. infestans*. In *P. infestans* populations, *Avr3a* has two major allelic variants encoding the proteins AVR3a^KlY147del^and AVR3a^EM^, which differ in two amino acids in the mature protein and are differentially perceived by the potato immune receptor R3a^63–67^. Contrary to AVR3a^EM^, AVR3a^KI^ induces R3a-mediated resistance and confers avirulence to homozygous or heterozygous strains of the pathogen^63^. In host plants that do not carry R3a, AVR3a^KI^ effectively suppresses the cell death induced by *P. infestans* INF1 elicitin and is thought to contribute to pathogen virulence through this and other immune suppression activities^64, 65, 68^. Remarkably, AVR3a, a mutant with a deleted C-terminal tyrosine residue, is not affected in activation of R3a but fails to suppress INF1-cell death demonstrating that distinct amino acids condition the two AVR3a activities^65^. Moreover, AVR3a^KI-Y147del^ cannot bind or stabilize the plant E3 ubiquitin ligase protein CMPG1, which is required for INF1-cell death, further uncoupling the effector activities^65, 67^. The current model is that AVR3a, but not AVR3a^KI-Y147del^, binds and stabilizes CMPG1 to suppress BAK1/SERK3-regulated immunity triggered by INF1 during the biotrophic phase of *P. infestans* infection^67^.

This study was prompted by our discovery that natural variants of the *P. infestans* effector AVR3a (AVR3a^KI^ and AVR3a^EM^) and the AVR3a^KI-Y147del^ mutant suppress FLS2-dependent early responses. This contrasts sharply with the differential activities of these three AVR3a variants in suppressing INF1 cell death, another BAK1/SERK3 dependentpathway. TheabilityoftheAVR3amutantto suppress FLS2-dependent responses revealed that AVR3a can suppress BAK1/SERK3-dependent responses in a CMPG1-independent manner. Furthermore, AVR3a reduced the internalization of the activated FLS2 receptor but did not interfere with its nonactivated plasma membrane localization indicating that this effector might target cellular trafficking initiated at the cell periphery. Consistent with this model, we found that AVR3a associates with a plant GTPase dynamin (NtDRP2) involved in receptor-mediated endocytosis implicating AVR3a in a cellular trafficking complex. Further, we found that NbDRP2 is required for internalization of FLS2, and that this protein plays a role in modulating the activity of immune receptors. Overexpression of NtDRP2-1 suppressed PRR-dependent accumulation of reactive oxygen species (ROS). We conclude that AVR3a associates with a key cellular trafficking and membrane-remodeling complex required for PRR endocytic trafficking.

## RESULTS

### AVR3a suppresses PTI in a CMPG1-independent manner

To determine the degree to which AVR3a suppresses PAMP-elicited defense responses besides INF1-elicited immunity, we measured the oxidative burst production and defense gene induction in *N. benthamiana* triggered by bacterial and oomycete elicitors. Plants transiently expressing epitope tagged variants of AVR3a^KI-Y147del^ (FLAG-AVR3a^KI^, FLAG-AVR3a^EM^, FLAG-AVR3a^KI-Y147del^) or the vector control (pBinplus: GFP) were treated with flg22 (100 nM) or INF1 (10 µg/ml) and the transient accumulation of reactive oxygen species (ROS) was followed over 45 minutes or 22 hours, respectively. We observed that all variants of AVR3a reproducibly suppressed flg22-induced ROS to the same extent whereas AVR3a^KI^ suppressed INF1-induced ROS more effectively than the other two variants consistent with previous reports^64, 65^ (Figure 1A, C). We observed similar results after flg22 treatment in *N. benthamiana* and *A. thaliana* plants stably expressing AVR3a validating the transient expression assays (Figure S1A, B).

**Figure 1.**
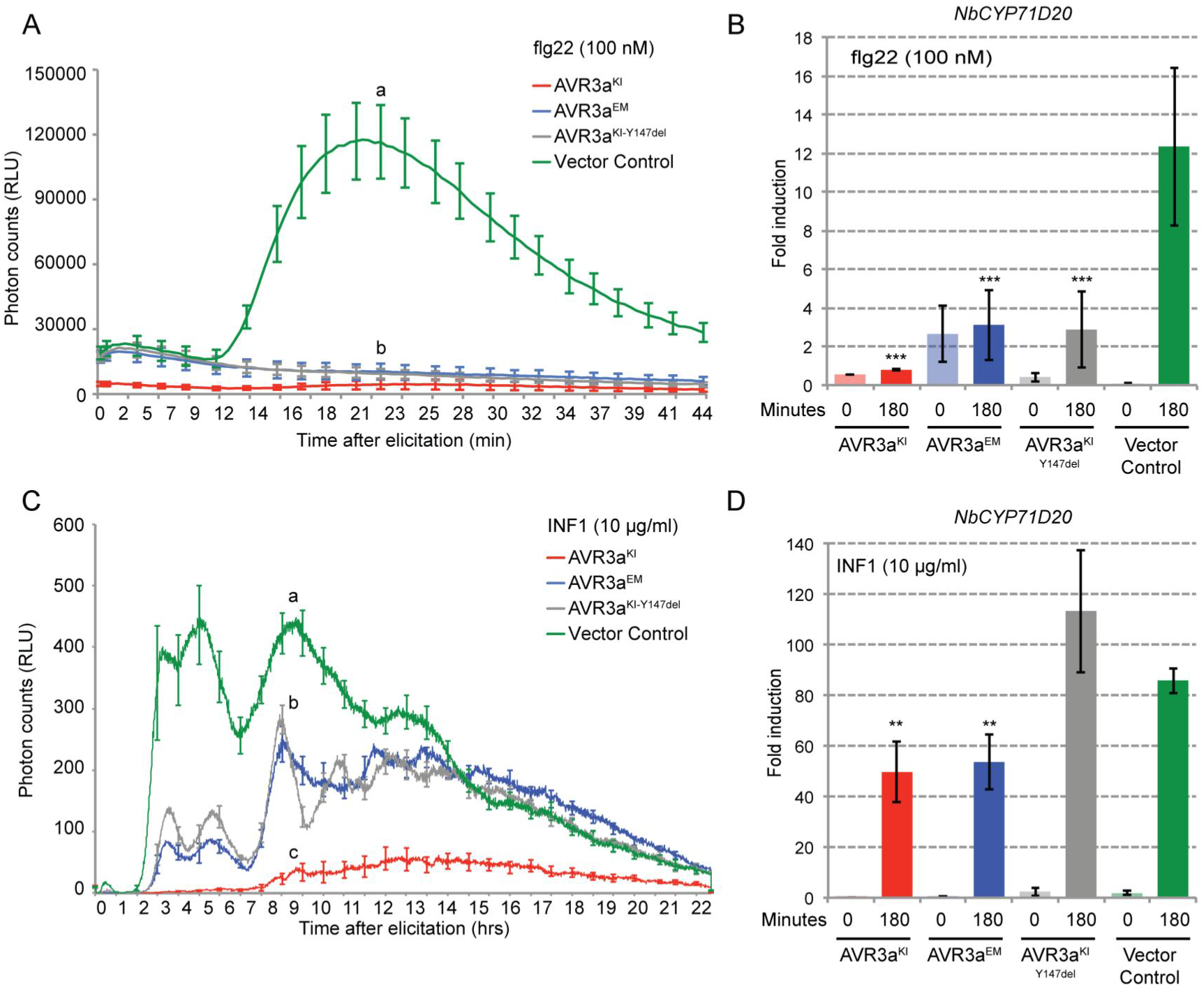
AVR3a suppresses elicitor-induced reactive oxygen species (ROS)production and gene induction. (A, C) Time course experiments measuring ROS production in relative light units (RLU) in response to 100 nM flg22 (A) or 10 µg/ml INF1[Pi] (C) in *N. benthamiana* transiently expressing FLAG-AVR3a^KI^ (red), FLAG-AVR3a^EM^ (blue), FLAG-AVR3a^KI-Y147del^ (grey) or Vector Control (ΔGFP, green). Different letters above the graph indicate significant differences at *P* < 0.001 assessed by one-way ANOVA followed by TukeyHSD test. (B, D) Quantitative RT-PCR analysis of expression of the marker gene *NbCYP71D20* by 100 nM flg22 (B) or 10 µg/ml INF1[Pi] (D) in *N. benthamiana* stably expressing the same FLAG-AVR3a constructs. Marker gene expression was assessed at time 0 and 180 minutes after elicitor treatment and transcript levels were normalized to the *NbEF1α* housekeeping gene. Results are average SE (n = 3 technical replicates). Statistical significance was assessed by one-way ANOVA followed by TukeyHSD test. \*\**P* < 0.01; \*\*\**P* < 0.001. Similar results were observed in at least three independent experiments.

The immune responses triggered by another bacterial PAMP, EF-Tu, overlap and share signaling components with those triggered by flagellin^28, 69, 70^. Therefore, we tested the effect of AVR3a on ROS triggered by the EF-Tu derived peptide elf18. We found that elf18 ROS production was significantly impaired to a similar extent by all variants of AVR3a (Figure S2A). In contrast, the BAK1/SERK3 independent ROS response to chitin was not affected by any of the AVR3a variants (Figure S2B). These results suggest that AVR3a suppresses BAK1/SERK3-dependent pathways but not BAK1-independent immune responses.

**Figure 2.**
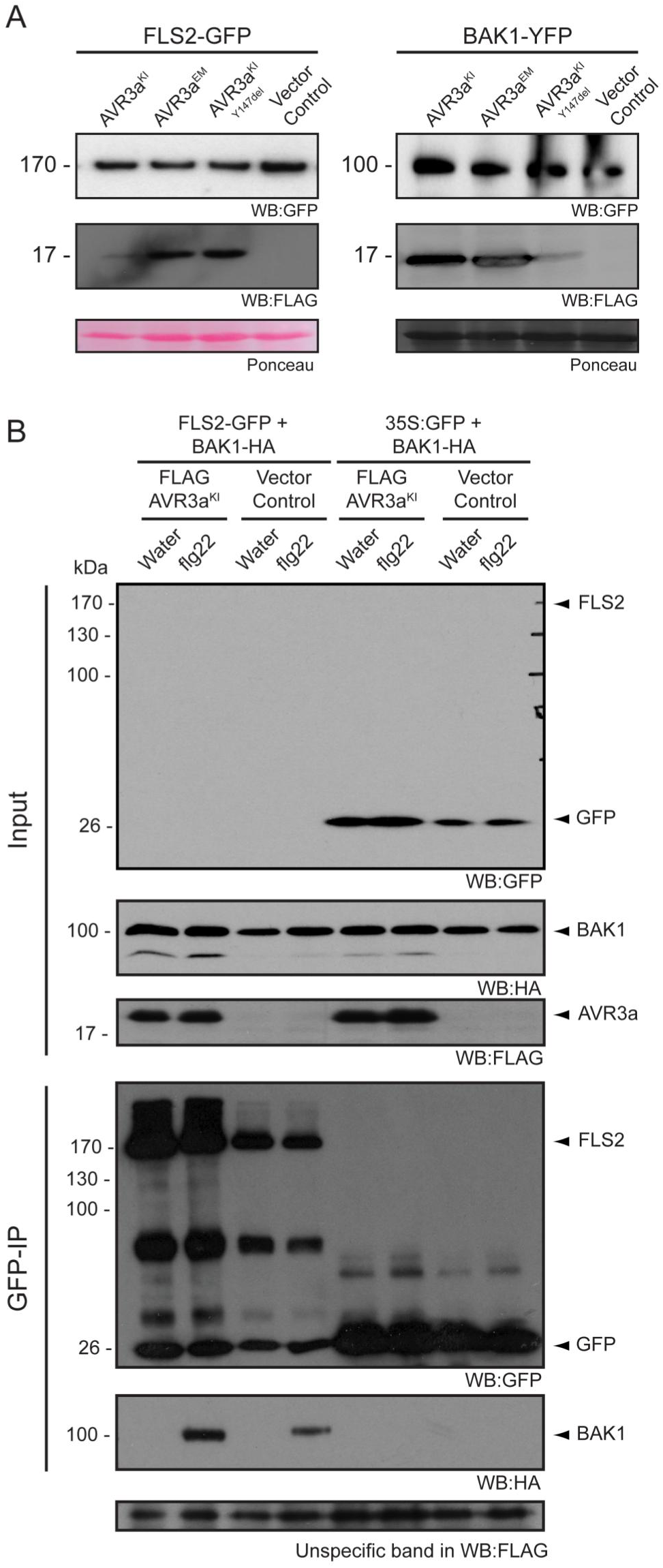
AVR3a does not affect protein accumulation of the nonactivated receptors FLS2 and BAK1 or their association *in planta*. (A) Transient co-expression in *N.benthamiana* of FLS2-GFP or BAK1-YFP with FLAG-AVR3a^KI^or FLAG-AVR3a^EM^or FLAG-AVR3a^KI-Y147del^ or Vector Control (ΔGFP) as indicated. Total protein was extracted at 2.5 days post infiltration (dpi) with GTEN and 80% of the total extract was subjected to immunoprecipitation with 8 µl of anti-GFP agarose beads (Chromotek) to enrich for the GFP-tagged proteins and detected using anti-GFP antibody. Equal amounts of protein were analyzed (Ponceau lane) in all cases. (B) Transgenic *N. benthamiana* expressing FLAG-AVR3a^KI^ or Vector Control (ΔGFP) were transiently infiltrated with a mix (1:1) of FLS2-GFP/BAK1-HA and treated with flg22 (100 nM) or water for 15 minutes. Treated leaf-tissue was collected at 2.5 dpi and subjected to co-immunoprecipitation with 25 µl GFP agarose beads (Chromotek). Purified complex formation after flg22 treatment of FLS2-GFP and BAK1-HA was analyzed by immunoblotting with the specified antibodies. 35S: GFP/BAK1-HA was used as control. Note that FLS2-GFP was almost never detected in the total extracts (Input) but became visible by western blot analysis after immunoprecipitation.

One of the outcomes of PAMP elicitation is transcriptional reprogramming of the plant, a defense response downstream of ROS^9^. Therefore, we monitored the effect of AVR3a on gene expression of the previously characterized PTI marker genes *NbCYP71D20* and *NbACRE31* ^26,71^. *NbCYP71D20* expression was induced ~12-fold by flg22 and ~80-fold by INF1 in control plants (Figure 1B, D). All AVR3a variants reduced the induction of *NbCYP71D20* by flg22 by approximately 80% (Figure 1B) whereas reduction of INF1-elicited gene induction was about 40% for AVR3a^KI^ and AVR3a^EM^ with no reduction observed for the AVR3a^KI-Y147del^ variant (Figure 1D). Similarly, induction of *NbACRE31* by flg22 treatment decreased by 80% in the presence of all variants of AVR3a (Figure S1C). In contrast, AVR3a did not suppress *NbACRE31* expression after INF1 elicitation, although INF1 induction of this gene was very low (Figure S1D).

Overall, our results indicate that in addition to suppressing INF1 cell death^64, 65, 67^, AVR3a has the capacity of supressing PTI responses mediated by the BAK1/SERK3 dependent cell surface receptors FLS2 and EFR. Remarkably, all the variants of AVR3a^KI-Y147del^, including AVR3a, which neither suppresses INF1 cell death nor interacts with the E3 ligase CMPG1, suppressed flg22 responses to the same extent. These findings indicate that the newly identified suppression activity is CMPG1 independent and that the AVR3a effector may suppress PTI through multiple mechanisms.

### AVR3a does not alter receptor levels at the cell surface or receptor complex formation

The bacterial effector AvrPtoB targets FLS2 for degradation to suppress plant immunity^72^. This prompted us to determine whether the suppression of flg22-triggered responses by AVR3a involved perturbation of FLS2 or BAK1 protein accumulation or complex formation^73, 74^. To address this question, we transiently co-expressed AVR3a variants with FLS2-GFP or BAK1/SERK3-YFP in *N. benthamiana* and assessed the fusion protein levels. We found that AVR3a did not alter FLS2 or BAK1/SERK3 protein accumulation *in planta* (Fig 2A). Next, we tested whether AVR3a^KI^ perturbed heterodimerization of FLS2 with BAK1/SERK3 after flg22 treatment^25, 26, 28^. We used leaves of transgenic *N. benthamiana* plants stably expressing AVR3a^KI^ or a vector control and transiently expressed FLS2-GFP and BAK1/SERK3-HA. In the presence of AVR3a^KI^ a double band appeared for BAK1/SERK3 total protein extracts (Figure 2B, WB: HA) that was not seen after immunoprecipitation. However, this observation was not consistent between experiments and probably is the result of protein degradation during protein extraction. After co-immunoprecipitation of FLS2 and BAK1/SERK3 we saw no consistent effect of AVR3a on flg22-mediated recruitment of BAK1/SERK3 into the FLS2 signaling complex (Figure 2B). In summary, these results suggest that AVR3a does not interfere with receptor protein accumulation or complex formation and that the effector suppression of flg22 responses most likely occurs downstream of FLS2/BAK1 heterodimerization.

### AVR3a interferes with FLS2 internalization

We hypothesized that AVR3a alters the subcellular distribution of FLS2 and/or BAK1/SERK3 to perturb their activities. To determine the effect of AVR3a on subcellular distribution of the receptors, we transiently co-expressed FLS2-GFP or BAK1/SERK3-YFP with AVR3a variants in *N. benthamiana* and assessed the localization of the inactive receptors by confocal microscopy. In both cases, the plasma membrane localization of FLS2 and BAK1/SERK3 remained unaltered in the presence of AVR3a^31, 42^ (Figure 3A). It is well established that FLS2 activation after flg22 perception leads to transient endocytosis and accumulation of the receptor in mobile endosomal compartments^41, 42, 44^. Therefore, we tested whether AVR3a has an effect on the subcellular distribution of the activated receptor. Similar to the observations by Choi et al.^44^, FLS2 internalization was detected at 80 minutes to 140 minutes post elicitation in *N. benthamiana* leaves (Figure 3B). Remarkably, FLS2 endosomal localization was partially inhibited in plants expressing AVR3a^KI^ (Figure 3B). To quantify this observation, we counted the number of images that showed a distinct fluorescent signal in punctate structures reminiscent of endosomes after treatment with flg22 or water. Vesicle-like structures were detected in less than 5% of the cells in the control treatment, whereas about 80% of the flg22-treated cells showed fluorescence in vesicle-like structures (Figure 3B, 3C). In contrast, only 40% of the cells expressing AVR3a and treated with flg22, showed endosomal localization of FLS2 (Figure 3C).

**Figure 3.**
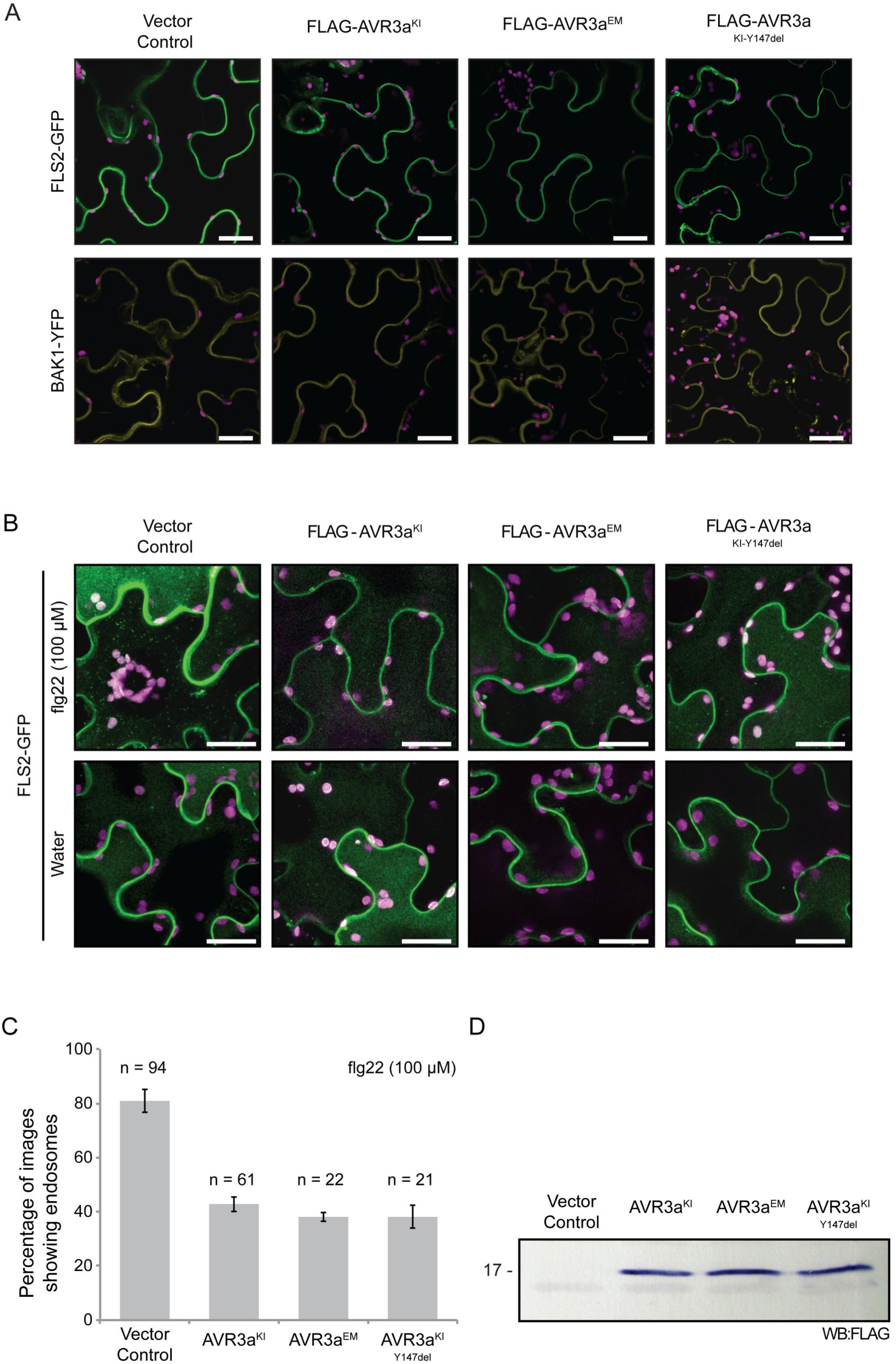
AVR3a partially suppresses the endosomal localization of the activated FLS2 receptor. (A) Confocal microscopy in *N. benthamiana* epidermal leaf cells stably expressing FLAG-AVR3a^KI^, FLAG-AVR3a^EM^, FLAG-AVR3a^KI-Y147del^ or Vector Control (ΔGFP) and infiltrated with FLS2-GFP or BAK1-YFP. The effector AVR3a did not alter the nonactivated FLS2 (green) or BAK1 (yellow) plasma membrane subcellular localization at 2.5 days post infiltration (dpi). Bar = 25 µm. Plastids autofluorescence (purple) is shown. (B) Standard confocal images of FLS2-GFP in *N.benthamiana* epidermal leaf cells under the same conditions described above,challenged for 90 to 120 minutes with 100 µM flg22 or water as indicated. Representative images show a clear accumulation of FLS2-GFP in intracellular vesicles (green dots) specifically after flg22 elicitation in the vector control plants whereas this distinct re-distribution of FLS2-GFP was partially inhibited by the presence of all variants of AVR3a. All images are a maximum projection of 20 slices taken at 1-µm intervals. Same confocal settings were used to acquire all images. Bar = 25 µm. Plastids auto-fluorescence (purple) is shown. (C) Quantification of the effect of AVR3a on FLS2-GFP endocytosis. Data was collected from 7 independent experiments showing similar results. The histogram depicts the number of images that showed FLS2-GFP endosomes as a percentage of the total number of images analyzed as indicated. Error bars are SE values. (D) Western blot probed with anti-FLAG antibody showing the expression of AVR3a in transgenic *N. benthamiana* plants.

We then examined whether AVR3a perturbs the subcellular localization of other cell surface receptors. First, AVR3a did not result in any changes in the membrane localization of the inactive immune receptors EFR and CERK1 (Figure S3A). Next, we determined whether inhibition of receptor internalization is specific to FLS2 or a general interference with the endocytic process. For this we used BRI1, a receptor involved in development that also requires BAK1/SERK3 for its activity and shows constitutive internalization^32^. Using the same experimental procedure described above, we observed that BRI1-GFP fluorescent signal from vesicle-like structures was unaltered in the presence of AVR3a^KI^ (Figure S4A, S4B). These results indicate that AVR3a targets FLS2 or a host component required for FLS2 internalization to modulate early defense responses without generally affecting receptor mediated endocytosis.

**Figure 4.**
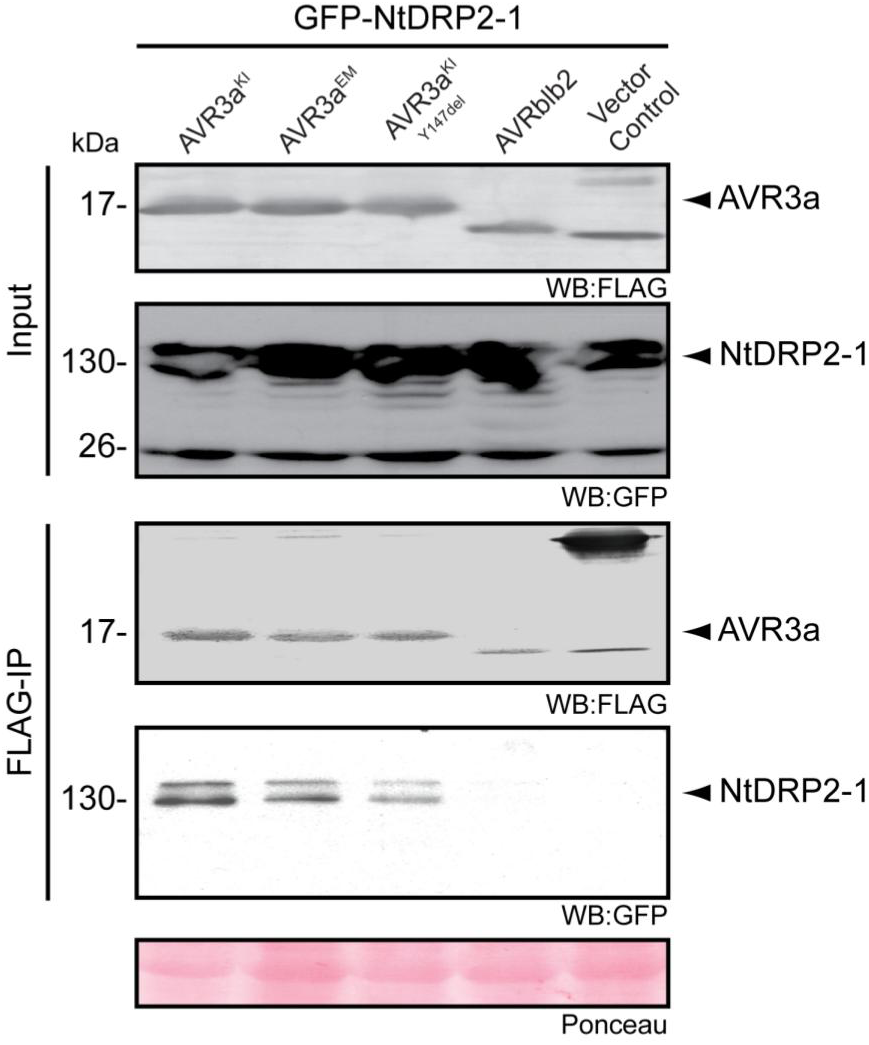
AVR3a associates with NtDRP2-1*in planta*. GFP-NtDRP2-1 wastransiently co-expressed with FLAG-AVR3a^KI^ or FLAG-AVR3a^EM^ or FLAG-AVR3a^KI-Y147del^ or FLAG-AVRblb2 or Vector control (FLAG-RFP) in *N. benthamiana*. Immunoprecipitates obtained with anti-FLAG beads (SIGMA) and total protein extracts were immunoblotted with the appropriate antisera. Protein was extracted 2.5 days post infiltration. Experiment repeated at least three times with similar results.

### AVR3a associates with a plant protein involved in cellular trafficking

AVR3a was previously shown to bind the E3 ligase CMPG1^67^. To determine what additional host proteins AVR3a associates with, we used co-immunoprecipitation of FLAG-AVR3a^KI^ in *N. benthamiana* followed by liquid chromatography-tandem mass spectrometry (LC-MS/MS) analysis using previously described methods^75^. We identified four proteins that associated with AVR3a^KI^ that had unique peptide counts compared to the controls in two biological replicates (Table S1). We selected a homolog of the GTPase dynamin-related protein (DRP) (NCBE_074039.1) for further studies based on the established role of dynamins in cellular trafficking and clathrin-mediated endocytosis in animal systems^76^. In mammals, dynamin is a five-domain protein consisting of a GTPase domain, an unstructured middle domain, a Pleckstrin-homology domain, a GTPase effector (GED) domain and a proline-rich domain, and its functions range from vesicle scission during endocytosis to the formation of the tubular-vesicular network during cytokinesis^76^. In Arabidopsis, DRP2A and DRP2B (previously known as ADL6 and ADL3, respectively) are the only dynamin-related proteins (DRP) that have similar domain architecture to other canonical dynamins^76–79^. Database searches using tBlastN and NCBE_074039.1 as a query revealed that DRP2A and DRP2B were the closest homologs for NCBE_074039.1. Therefore, we designed primers based on the Arabidopsis DRP2A and DRP2B sequences and the partial *N. benthamiana* sequences available at the time of the experiment. However, these primers did not result in any amplicon from *N. benthamiana* cDNA but we cloned two putative DRP2 proteins from *N. tabacum*, termed NtDRP2-1 and NtDRP2-2, which are 99% similar to each other. Sequence analysis revealed that NtDRP2-1 and NtDRP2-2 have the classical five-domain structure of canonical dynamin proteins^76^ and have 77% amino acid similarity to the Arabidopsis DRP2 proteins (Figure S5). To determine the evolutionary relationship of NtDRP2-1/2 to other plant dynamin-like proteins, we first searched the genomes of solanaceous species, Arabidopsis and other dicot plants for proteins containing the dynamin signature^80^. Next, we constructed a maximum likelihood tree based on the amino acid sequence alignment of the conserved GTPase-domain of 45 identified dynamin related proteins (Figure S6, Table S2). The resulting tree confirmed that NtDRP2-1/2 are most related to AtDRP2A and AtDRP2B among the 11 Arabidopsis DRPs (Figure S6).

We then investigated the subcellular localization of NtDRP2 to determine whether it is consistent with the presumed function of DRPs in cellular trafficking. Transient expression of GFP-NtDRP2-1 and GFP-NtDRP2-2 in *N. benthamiana* followed by confocal microscopy revealed that GFP-NtDRP2-1 and GFP-NtDRP2-2 mainly localized at the plasma membrane as seen by the formation of thin cytoplasmic strands at points of adhesion between the plasma membrane and the cell wall (Hechtian strands) (Figure S7B, S7C). In addition, GFP-NtDRP2-1 and GFP-NtDRP2-2 localized to the cytosol in a punctate pattern (Figure S7C). This subcellular distribution is consistent with the reported localization of classical dynamins in animal, yeast and Arabidopsis^76,77,81,82^.

Next we validated the association of AVR3a with the cloned NtDRP2 *in planta* by co-immunoprecipitation analysis. All three AVR3a variants, including AVR3a^KI-Y147del^, co-immunoprecipitated with GFP-NtDRP2-1 when expressed in *N. benthamiana* as FLAG epitope tagged proteins (Figure 4). The RXLR effector AVRblb2 was used as a negative control in these experiments and did not co-immunoprecipitate with NtDRP2-1 (Figure 4). We also used the *Phytophthora capsici* effector PcAVR3a-4^61^ as an additional negative control because this AVR3a homolog does not suppress the ROS burst induced by flg22 (Figure S8B). Indeed, in side-by-side co-immunoprecipitation experiments there was only a weak NtDRP2-1 band with PcAVR3a-4 that only became visible with increased exposure times (Figure S8A). Overall, these results confirm that all AVR3a variants associate with NtDRP2-1, and suggest this association may be linked to the ability of AVR3a to suppress flg22-responses.

### NtDRP2 dynamin is required for FLS2 internalization

To further investigate the link between AVR3a effector activities and NtDRP2 dynamin, we investigated the degree to which this dynamin is required for FLS2 internalization using RNAi silencing experiments. We designed silencing constructs that target *N. benthamiana* DRP2 *Nb05397* (*NbDRP2-1*) and *Nb31648* (*NbDRP2-2*), which are the most closely related *N. benthamiana* DRP2 to the tobacco NtDRP2-1 and NtDRP2-2 (Figure S6). Reports in Arabidopsis showed that members of the DRP2 family are implicated in cell cytokinesis and post-Golgi vesicular trafficking and are essential for development with *drp2* double mutants exhibiting pleiotropic developmental defects^77–79, 83^. In addition, in an independent study, Smith and colleagues^84^ recently showed that DRP2B functions in flg22-signaling and bacterial immunity in Arabidopsis^84^. Using virus-induced gene silencing (VIGS) in *N.benthamiana*, we observed that plants silenced for *NbDRP2-1* and *NbDRP2-*2 displayed severe developmental defects and ultimately become necrotic and died confirming the essential role of dynamins in plant development (Figure S9). To circumvent VIGS-induced lethality, we transiently expressed a *DRP2*-targeted hairpin-silencing construct in fully developed *N. benthamiana* leaves to silence *Nb05397* and *Nb31648* (Figure S10A). Using quantitative RT-PCR, we determined that the hairpin construct significantly reduced the mRNA levels of *Nb05397* and *Nb31648* but not of the closely related *N. benthamiana DRP2* genes *Nb11538* and *Nb09838*, indicating that the RNAi silencing is specific to the targeted *N. benthamiana DRP2* genes (Figure S10B). We also confirmed that the silenced epidermal cells remained viable by staining with propidium iodide, which is excluded from cells with intact plasma membranes (Figure S10C).

Next, we used the hairpin RNAi system to determine the effect of silencing *NbDRP2* in *N. benthamiana* leaves on flg22-induced internalization of FLS2-GFP. We found that FLS2-GFP containing vesicles were absent in the *NbDRP2* silenced leaves upon flg22 treatment, but were clearly visible in control silenced leaves (Figure 5A, 5C). Moreover, silencing *NbDRP2* did not alter accumulation of FLS2 at the plasma membrane in water treated samples (Figure 5A), indicating that DRP2 only affects FLS2 after activation. To determine whether the effect of *NbDRP2*-silencing on receptor internalization is specific to FLS2, we performed the same *NbDRP2* RNAi experiments with the brassinosteroid receptor BRI1. Remarkably, BRI1-GFP constitutive endocytosis was not affected in leaves silenced for *NbDRP2* and a similar number of BRI1-GFP labelled vesicles was observed in the *NbDRP2* RNAi treatment compared to the negative control (Figure S4C, S4E). We conclude that *NbDRP2* is essential for ligand-induced endocytosis of FLS2, and that this requirement might be specific for immune-related receptors. Importantly, these findings are consistent with our earlier finding that AVR3a interferes with the internalization of the activated FLS2 receptor but not BRI1 (Figure 3B, 3C and Figure S4A, S4B).

**Figure 5.**
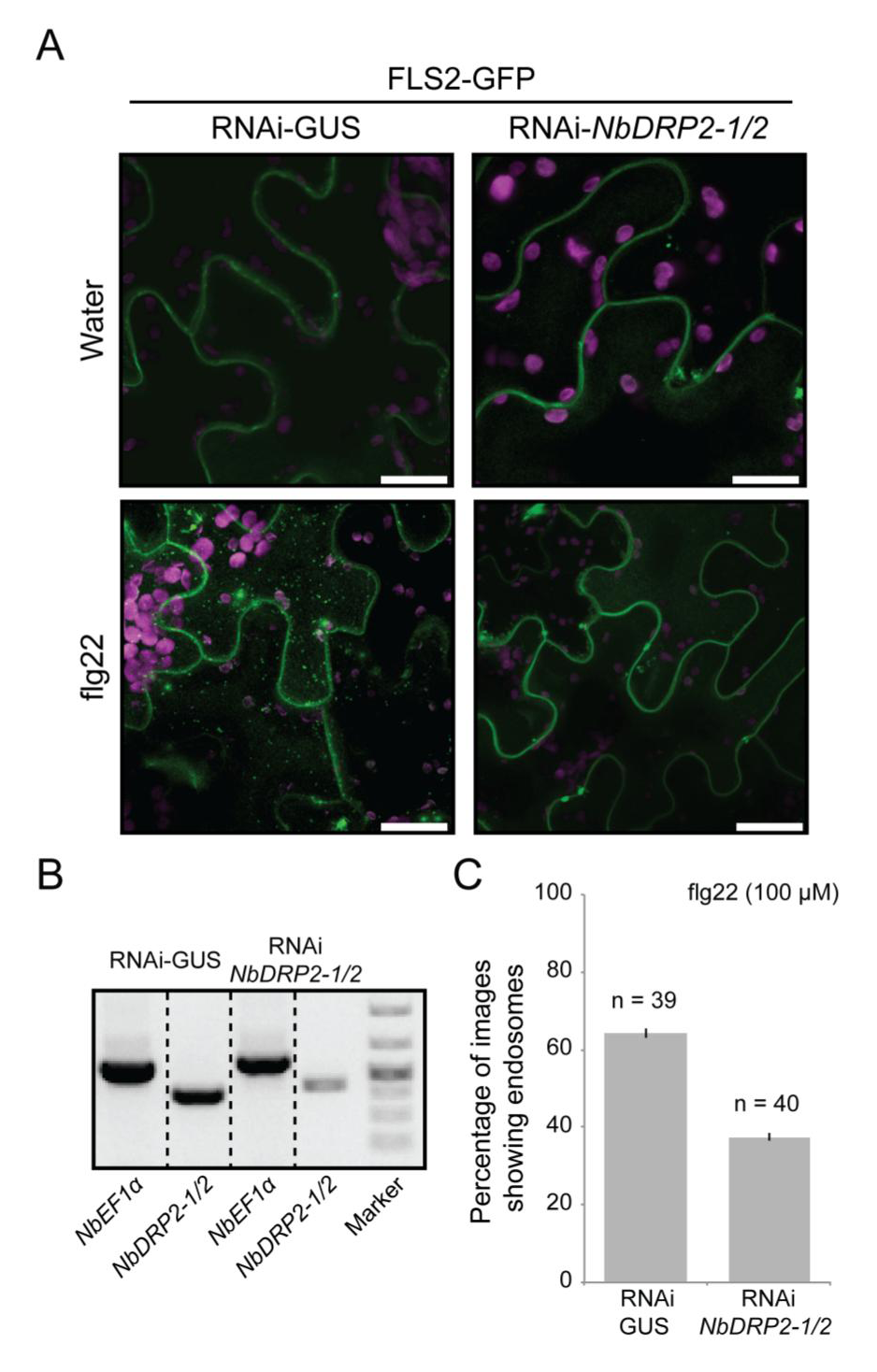
*NbDRP2-1/2*is partially required for FLS2 internalization. (A) Confocalimaging in *N. benthamiana* epidermal cells reveals that decreased *NbDRP2-1/2* expression leads to a significant reduction of subcellular distribution of FLS2-GFP in endosomes (green dots) after flg22 elicitation (100 µM). *N. benthamiana* leaves were transiently expressing FLS2-GFP plus a hairpin-silencing construct RNAi-*NbDRP2-1/2* or the silencing vector control RNAi-GUS. Imaging was done at 2.5 days postinfiltration. All images are a maximum projection of 21 slices taken at 1-µm intervals. Same confocal settings were used to acquire all images. Bar = 25 µm. Plastids autofluorescence (purple) is shown. (B) Validation of RNAi-*NbDRP2-1/2* silencing by RT-PCR in leaf-discs collected from the same leaves used for microscopy. Bands were grouped in the figure and dashed black lines represent different parts of the same gel. (c)Quantification of the effect of RNAi-*NbDRP2-1/2* on FLS2-GFP endocytosis. Data was collected from 6 independent experiments showing similar results. The histogram depicts the number of images that showed FLS2-GFP endosomes as a percentage of the total number of images analyzed as indicated. Error bars are the SE values.

### Overexpression of NtDRP2 dynamin suppresses flg22-induced ROS burst

To assess the role of Solanaceous DRP2 in early defense responses, we first determined the degree to which silencing of *NbDRP2* affects flg22-induced ROS production. These experiments were inconclusive as we failed to observe consistent effects on ROS burst in *NbDRP2*-silenced *N. benthamiana* leaves (data not shown). We then determined the effect of overexpressing NtDRP2 in *N. benthamiana* on ROS production. Consistent with the finding that AtDRP2B is a negative regulator of flg22-triggered ROS^84^, we found that NtDRP2-1/2 decreased flg22-induced ROS burst by more than 50% compared to control plants (Figure 6A). In addition, NtDRP2-1 significantly reduced the ROS production in response to INF1 and chitin treatments (Figure 6B, 6C). These results indicate that the effect of DRP2 on ROS responses is not limited to BAK/SERK3-dependent pathways and that NtDRP2 dynamins modulate early signaling responses induced by several PAMPs.

**Figure 6.**
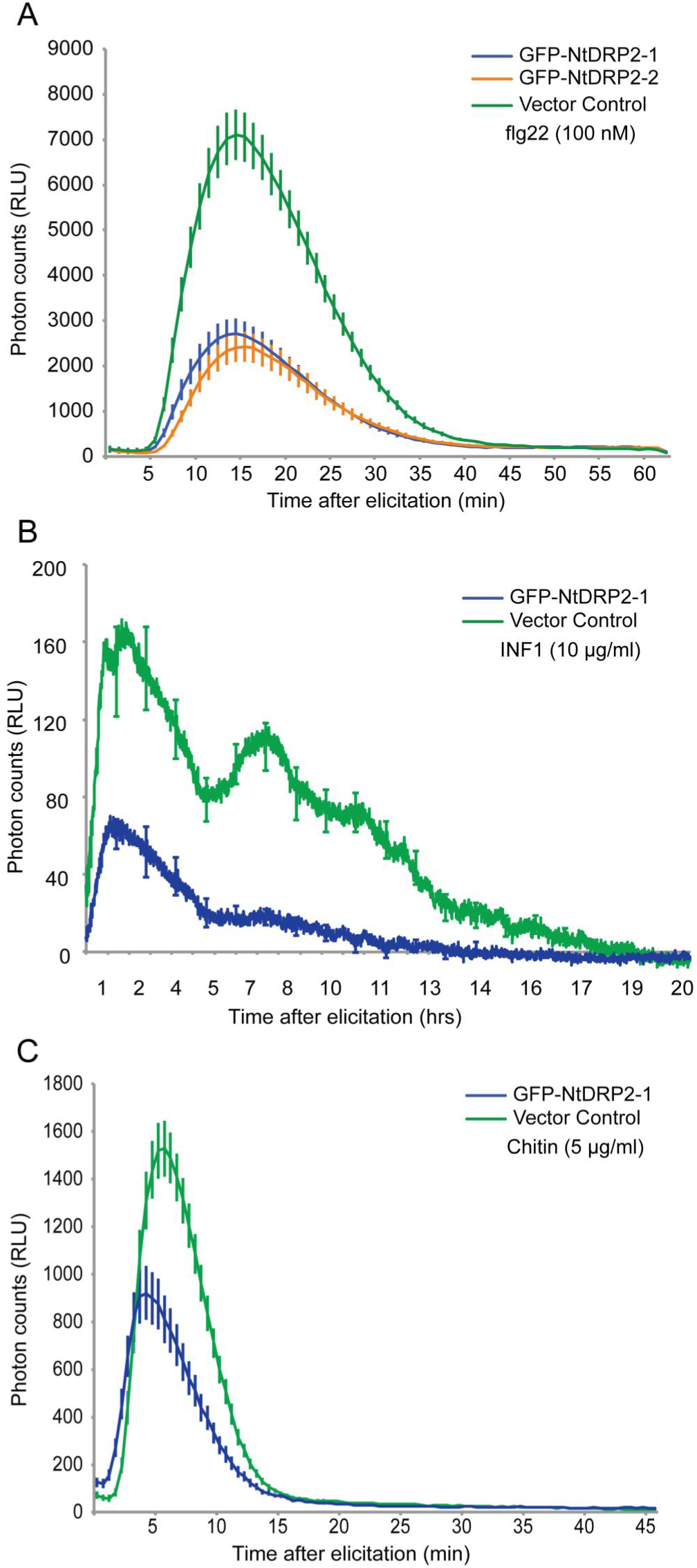
Overexpression of NtDRP2 reduces the accumulation of ROS upon PAMP perception. (A, B, C) *Agrobacterium tumefaciens*-mediated transient expression of GFP-NtDRP2-1 or Vector Control (35: GFP) in *N. benthamiana* leaves showed that ROS production after elicitation with flg22 (A), INF1 (B) or chitin (C) is reduced in the presence of NtDPR2. Total ROS production was measured in relative light units (RLU) over time after treatment with 100 nM flg22 (A), with 10 µg/ml INF1[Pi] (B) or with 5 µg/ml chitin (SIGMA) (C). ROS was measured at 3 days post infiltration. Similar results were observed in at least four independent experiments.

## DISCUSSION

Even though perception of microbe-associated molecular patterns significantly contributes to basal defense of plants against the Irish potato famine pathogen *P.infestans*, our knowledge of how effectors subvert basal immunity is patchy. One of the best studied effectors of *P. infestans* is the RXLR-WY type effector AVR3a^63–68, 85^. In this study, we further characterized the virulence activities of AVR3a and discovered that AVR3a suppresses early defense responses mediated by the cell surface immune modulator BAK1/SERK3, which contributes to basal immunity against *P. infestans*^36^. More specifically, we discovered that AVR3a reduces internalization of the activated pattern-recognition receptor FLS2 but does not interfere with the plasma membrane localization of nonactivated FLS2 nor does it perturb the steady state levels of this immune receptor or other PRRs. Furthermore, we found that AVR3a associates with DRP2, a *Solanaceae* member of the plant GTPase dynamin family that mediate endocytosis and membrane remodeling^86^. Interestingly, DRP2 is required for FLS2 internalization and DRP2 overexpression reduces the ROS burst induced by the flg22 flagellin peptide. Our results indicate that AVR3a associates with a key cellular trafficking and membrane-remodeling complex, and that this DRP2 dynamin complex is involved in immune receptor-mediated endocytosis and signaling.

Several bacterial type III secretion system effectors target cell surface immune receptor complexes to modulate their activities^13,72,74^. AVR3a is an example of a filamentous plant pathogen effector that has evolved to deregulate plant immune signaling^64, 67, 68^. Our work is consistent with the finding that AVR3a blocks signal transduction cascades initiated at the plasma membrane after pathogen perception^68^. However, interference with FLS2 signaling and endocytosis does not involve the interaction with the E3 ligase CMPG1 since AVR3a^KI-Y147del^, a variant that is unable to bind and stabilize CMPG1, is still able to suppress FLS2 ROS burst and endocytosis (Figure 1A; Figure 3B, 3C). Therefore, AVR3a is a multifunctional effector that can suppress BAK1/SERK3 mediated immunity through at least two different pathways, possibly by acting at different sites or at different time points during immune signaling.

The step at which AVR3a interferes with FLS2 signaling is unclear. AVR3a is unlikely to completely inhibit DRP2 activity since DRP2 silencing resulted in plant lethality unlike transient or stable *in planta* expression of AVR3a. AVR3a did not alter subcellular localization of FLS2 or BAK1/SERK3 (Figure 3A) and it did not affect ligand induced complex formation between FLS2 and BAK1/SERK3 (Figure 2). However, the receptor complex somehow remained inactive in the presence of AVR3a given that downstream signaling cascades were not activated. Given that elicitor-induced internalization of FLS2 occurs in a BAK1-dependent manner^42, 43^ it is possible that the effect of AVR3a in FLS2 internalization occurs via BAK1/SERK3. However, BAK1/SERK3 C-terminal fusions are partially impaired in early PTI responses^87^ making it difficult to assess the extent to which BAK1/SERK3 undergoes endocytosis after flg22 perception. Interestingly, AVR3a exhibits some degree of specificity in suppressing PAMP-triggered immunity. Unlike the *P. syringae* effector AvrPtoB, which blocks the responses elicited by bacterial flagellin and chitin^88–90^, none of the AVR3a variants suppressed chitin-induced ROS burst suggesting that AVR3a specifically suppresses BAK1/SERK3-dependent immune responses. At this stage, we propose that AVR3a targets BAK1/SERK3 cellular trafficking initiated at the cell periphery but additional work is needed to clarify the underlying mechanisms.

Our finding that AVR3a co-immunoprecipitates with DRP2 does not necessarily imply that these two proteins directly bind *in-planta* nor that DRP2 is a target of the effector. DRP2 could be a helper of AVR3a that functions as a cofactor or enables localization of AVR3a to particular subcellular compartments as defined by Win et al.^4^. Nonetheless, our results place AVR3a in proximity to a membrane-remodeling complex that is implicated in endocytosis of the cell surface pattern-recognition receptor FLS2. Remarkably, AVR3a affected FLS2 but not BRI1 endocytosis, indicating that the mechanisms for FLS2 and BRI1 internalization may be different. This is not completely unexpected, as previous reports have shown that the regulatory role of BAK1/SERK3 in BR and PTI signaling are distinct and can be mechanistically uncoupled^70, 87^. Moreover, there are different pools of BAK1/SERK3 that are not interchangeable between BRI1 and FLS2 and activation by BR or flg22 does not cross activate these signaling pathways^91^. Therefore, one possibility is that FLS2 and BRI1 share a common endocytic pathway but AVR3a only blocks the FLS2 receptor complex activation and prevents the downstream endocytosis. However, consistent with the specific activity of AVR3a on FLS2 internalization, DRP2 is specifically required for FLS2-endocytosis but not BRI1. This supports a model in which different internalization pathways for receptor endocytosis may occur following flg22 and brassinosteroid perception. For instance, AVR3a may perturb DRP2 functions only at sites where the activated PRR complexes accumulate and prevent receptor internalization. Our results indicate that BAK1/SERK3-associated membrane-bound receptors may be initially internalized via different pathways, which later on can converge at the late endosomal compartments.

In animal systems, canonical dynamin proteins mediate pinching off clathrin-coated vesicles from the membrane during constitutive endocytosis^52, 55^. In Arabidopsis, the two canonical dynamins DRP2A and DRP2B have been shown to play a role in endocytosis and to be genetically and functionally redundant^78, 83^. Recently, Smith et al^84^., showed that in Arabidopsis, FLS2 endocytosis is partially dependent on DRP2B, but not DRP2A^84^. Interestingly, the *Solanaceae* DRP2 family has expanded relative to Arabidopsis with no obvious orthologs of *DRP2A* and *DRP2B* (Figure S6). Consistent with Smith et al^84^., our results also indicate that the DRP2 family has evolved multiple activities, with the *Solanaceous* DRP2 being involved in FLS2 but not BRI1 internalization (Figure 5; Figure S4B). Possibly, plant DRPs have diversified to enable increased plasticity in response to biotic and abiotic stimuli. Further studies are clearly needed to fully understand the precise contributions of different plant DRPs to vesicle trafficking.

In summary, we link the *P. infestans* effector AVR3a to a membrane complex that includes the vesicle trafficking protein DRP2. This complex mediates endocytosis of the classic pattern recognition receptor FLS2 following activation by the flagellin-derived peptide flg22. Although the FLS2 co-receptor BAK1/SERK3 is required for basal immunity against the oomycete *P. infestans*, there is no evidence that FLS2, a receptor for bacterial flagellin, is activated and internalized during infection by this oomycete pathogen^39^. Therefore, AVR3a suppression of FLS2 response and endocytosis may indicate that this effector targets a common node shared by FLS2 and a yet to be discovered PRR involved in oomycete immunity. Future studies are required to address this possibility and further determine the various mechanisms by which the multifunctional RXLR-WY effector perturbs PTI signaling.

## MATERIALS AND METHODS

### Plant material

*N. benthamiana* and *A. thaliana* plants were grown and maintained under controlled environmental conditions at an average temperature of 23 - 25°C, with 45-65% humidity, and in long day conditions (16 hrs of light) throughout the experiments. *N.benthamiana* and *A. thaliana* were transformed^92^ with *A. tumefaciens* GV3101 or AGL1, respectively, carrying the following constructs: pBinplus: FLAG-AVR3a^KI^, pBinplus: FLAG-AVR3a^EM^, pBinplus: FLAG-AVR3a^KI-Y147del^, or pBinplus: GFP. Transformed plants were selected on Murashige-Skoog salts (MS) media containing selective antibiotic (kanamycin). Plates showing a segregation of 3:1 were selected and at least ten individual lines were selected to confirm expression of the transgene by western blot analysis with anti-FLAG antibody (SIGMA). Protein expression was confirmed for each individual plant once the plants reached the 3-week-old stage. Lines with different expression profiles were selected for further analysis. Transgenic lines used in this study were T4 and T3 for *N. benthamiana* and *A. thaliana* respectively.

### A. *tumefaciens-mediated transient gene expression assays in N. benthamiana*

*Agrobacterium tumefaciens* (strain GV3101) carrying the desired T-DNA construct was grown overnight at 28°C in Luria-Bertani culture medium with the appropriate antibiotics. Cells were harvested by centrifugation at 8000 g and resuspended in agro-infiltration media [5mM MES, 10 mM MgCl_2_, pH 5.6] prior to syringe infiltration into 3-4 week-old *N. benthamiana* plants. Each individual construct was infiltrated at a final OD_600mm_ of 0.3. Acetosyringone was added to the resuspended cultures at a final concentration of 150 µM and bacterial cultures were left at room temperature for 2 hours before infiltration.

## Silencing experiments

445-nucleotide cDNA fragment of *NbDRP2-1* (*Nb05397*, position 2100-2545 bp) was cloned into pTV00 (between SpeI/KpnI sites) using the following primers *NbDRP2-1*_SpeI, 5’ −GCGACTAGTATCAGCTCTAAAGGCGGTCA and *NbDRP2-*1_KpnI, 5’ −AAAAGGTACCGCTGTTGGGCTACTTTCTGC to form TRV2-*NbDRP2*. TRV2 plasmids were transformed into *A. tumefaciens* GV3101. For VIGS assays, *A.tumefaciens* containing TRV2 and TRV1 plasmids were grown and prepared individually for agro-infiltration assays as previously described^93, 94^. *A. tumefaciens* TRV1 and TRV2 were mixed in a 1:4 ratio to a final OD_600_ = 1.0 and infiltrated into 2-week-old *N. benthamiana* plants. Plants were analyzed and photographed three weeks after inoculation. TRV-ΔGFP was used as a negative control and TRV-SU (carrying a fragment from the phytoene desaturase gene) was used as a silencing control.

For transient silencing experiments, the same 445-nucleotide cDNA fragment of *NbDRP2-1* used for VIGS was cloned into pENTR/D-TOPO (Invitrogen) using the following primers: *NbDRP2-1*_hp_F2, 5’ – CACCATCAGCTCTAAAGGCGGTCA and *NbDRP2-1*_hp_R2, 5’ – GCTGTTGGGCTACTTTCTGC. The hairpin-silencing construct was generated by recombination into pHellesgate8 (Gateway LR recombination, Invitrogen) as described by Helliwell and Waterhouse^95^. The same procedure described above was used to generate the control silencing construct pHellsgate8-GUS (574 bp fragment size) using primers pH8_GUS_F1, 5’ – CACCCCAGGCAGTTTTAACGATCAG and pH8_GUS_R1, 5’ – GATTCACCACTTGCAAAGTCC. The final constructs were transformed into *A.tumefaciens* GV3101. Four-week-old *N. benthamiana* leaves were infiltrated with the hairpin silencing construct at a final OD_600_ = 0.3 either individually or co-expressed with a construct expressing the plant receptor under assessment. Further experiments (microscopy, analysis of gene silencing by qRT-PCR, PAMP elicitation and others described elsewhere in Materials and Methods) were performed three days after silencing.

## Reactive oxygen species measurement

Generation of reactive oxygen species (ROS) was measured as previously described^89^. Briefly, leaf-discs (16-24 per treatment, 4 mm diameter) from pre-infiltrated or transgenic *N. benthamiana* leaves were floated 16 hours in 200 µl of water in a 96-well plate. Solution was replaced by a luminol/peroxidase mix [17 mg/ml (w/v) luminol (Sigma); 10 mg/ml horseradish peroxidase (Sigma)] supplied with either 100 nM flg22 peptide (EzBiolab), 100 nM elf18 peptide (EzBiolab), 5 µg/ml chitin (Sigma), or 10 µg/ml purified INF1 [Pi] protein^36, 96^. Luminescence was measured over time (up to 1320 min) using an ICCD photon-counting camera (Photek, East Sussex, UK) and analyzed using company software and Microsoft Excel.

## Gene expression analysis

Transgenic *N.benthamiana* leaves expressing the construct sp Binplus: FLAG-AVR3a^KI^, pBinplus: FLAG-AVR3a^EM^, pBinplus: FLAG-AVR3a^KI-Y147del^ or pBinplus: GFP were treated for 0 and 180 minutes with flg22 (100 nM) or INF1[Pi] (10 µg/ml) on one side of the leaf and with Milli-Q water on the other side of the leaf as control. Total RNA was extracted using TRI reagent (Invitrogen) following manufacturer’s instructions. DNase treatment (Ambion) was performed according to manufacturer’s protocol and total RNA was quantified with a Nanodrop spectrophotometer (Thermo). 1.5 µg of DNase treated RNA was used for cDNA synthesis using SuperScript II reverse transcriptase (Invitrogen). qRT-PCR was performed with SYBR Green master mix (SIGMA) in triplicate per sample per gene. *NbEF1α* was used to normalize transcript abundance for the marker genes *NbACRE132*, *NbCYP71D20* and *NbACRE31*^71^. Primers used for amplification of *NbEF1α* and PAMP-induced marker genes in *N. benthamiana* have been reported previously^71^.

QRT-PCR analysis of *N. benthamiana* genes *Nb05397* (*NbDRP2-1*), *Nb31648* (*NbDRP2-2*), *Nb11538* and *Nb09838* was performed with SYBR Green master mix (SIGMA) in triplicate per sample per gene, using primers silDyn_97-48_F, 5’–CGATCGAGGAATTGACACAA and silDyn_97-48_R, 5’– GCCTGAGCAGCAGATATTACG to detect *Nb05397* and *Nb31648* or 38s_F4, 5’– TAATCGAGCAGCTGCTGTG and 38s_3’UTR_R, 5’-TAGGATCAAGCAGCAACTG to detect *Nb11538* and *Nb09838*. *NbEF1α*^71^ was used to normalize transcript abundance. 2.5 µg of DNase treated RNA was used for cDNA synthesis. Primers for *Nb11538* or *Nb09838* were predicted to specifically anneal at the 3’UTR of both sequences whereas primers for *Nb05397* and *Nb31648* were predicted to specifically anneal along the GTPase domain of *Nb05397* and *Nb31648*.

## Elicitor preparations

Chitin (crab shell chitin) and flg22 (QRLSTGSRINSAKDDAAGLQIA) peptides were purchased from SIGMA and EzBiolab, respectively and dissolved in ultrapure water. INF1 was purified from *P. infestans* 88069 by chromatography and the final working solution was dissolved in ultrapure water^36, 96^.

## Co-immunoprecipitation assays and western blot analysis

*N. benthamiana* leaves expressing pTRBO: AVR3a^KI^, pTRBO: AVR3a^EM^, pTRBO: AVR3a^KI-Y147del^, pTRBO: RFPorpTRBO: PcAVR3a-4with pK7WGF2: NtDRP2-1 or pK7WG2-3xHA: NtDRP2-1 constructs were harvested at 3 days post infiltration and ground in liquid N_2_ (see figure legends for more information). In Figure 2, pK7FWG2: FLS2 and pGWB14: BAK1 were co-infiltrated in *N.benthamiana* transgenic plants expressing pBinplus: FLAG-AVR3a^KI^ or pBinplus: GFP and treated with flg22 (100 nM) or water for 15 minutes. Protein extraction and immunoprecipitation was performed with 50 µ l of anti-FLAG resin (SIGMA) or 30 µ l GFP-affinity matrix (Chromotek) as previously described^28, 75^. Western blots analyses were performed as described by Oh et al.^97^,. Protein blotting was done with monoclonal FLAG primary antibody (SIGMA) and anti-mouse as secondary antibody (SIGMA), or monoclonal GFP-HRP antibody (Santa Cruz) or monoclonal HA-HRP antibody (Santa Cruz).

Sample preparation and liquid chromatography-tandem mass spectrometry (LC-MS/MS) analysis was performed as described in Caillaud et al^98^. Candidate AVR3a-associated proteins were identified from the peak lists searched on Mascot server v.2.4.1 (Matrix Science) against an in-house *N. benthamiana* database available upon request. We followed the same criteria in Caillaud et al^98^., for peptide identification. A complete list of all identified peptides for each AVR3a^KI^-associated protein is provided in Supplemental Table 3.

The pK7FWG2: FLS2 clone was generated from a pENTR: FLS2 and recombined into pK7FWG2 (GATEWAY, Invitrogen). The pGWB14: BAK1 clone is described in Roux et al., 2011 and Schwessinger et al., 2011.

## Cloning of *NtDRP2-1/2*

The AVR3a^KI^-associated protein NCBE_074039.1 sequence identified through the LC-MS/MS analysis was used as query to search the TAIR9 database. *Arabidopsisthaliana* proteins DRP2A and DRP2B were then used as query to perform a TBLASTN in the *Nicotiana benthamiana* draft genome scaffolds and tomato genome database (Solanum Genomic Network (SGN)). Partial sequences were used to design the primers Dyn_F3, 5’ – CACCATGGAAGCAATCGAGGAATTGGAGCAG and Dyn_R2, 5’ – TTATGATCTATAACCAGATCCAGACTGTGGTGG to amplify the open reading frame of the *Nicotiana tabacum* putative homologs of AtDRP2A/B from cDNA using Phusion proof reading polymerase (New England Biolabs). Amplicons were cloned into pENTR/D-TOPO (Invitrogen) and sequenced. Clones representing *NtDRP2-1* and *NtDRP2-2* were recombined into pK7WGF2, pK7WG2, and pK7WG2-3xHA to generate fusion proteins GFP-NtDRP2-1/2, non-tagged NtDRP2-1/2, and HA-NtDRP2-1/2, respectively. The final constructs were transformed into *A.tumefaciens* GV3101 and used for further analysis.

## Phylogenetic analysis

In Arabidopsis, GTPases with a PH domain belong to the DRP2 subfamily (Hong et al., 2003). Members of this subfamily, AtDRP2-A, AtDRP2-B, and the NtDRP2-1/2 cloned protein were used as queries to search for homologs of DRP2 in Solanaceous and other dicot plants. We performed TBLASTN searches against the *Nicotiana benthamiana* genome version 0.4.4, the tomato genome International Tomato Annotation Group release 2.3, the potato genome Potato Genome Sequencing Consortium DM 3.4 (solanaceous genomes available at http://solgenomics.net) and the NCBI database. To reveal the evolutionary relationship of these proteins, an alignment of the conserved GTPase domain was constructed using the multiple alignment software MUSCLE^99^ and a maximum likelihood (ML) tree was generated using the RAxML software^100^. To choose the best model for the ML tree, we estimated likelihoods of the trees constructed based on commonly used substitution models. The ML tree in Supplemental Figure 6 was constructed with GAMMA model of rate heterogeneity and Whelan and Goldman model (WAG), since the tree based on WAG showed the highest likelihood. We performed 500 non-parametric bootstrap inferences. Bootstrap values over 70% are shown. All Arabidopsis DRPs were included in the analysis to ensure proper estimation of the closest homologs of AtDRP2A/B. Sequences identifiers are available in Supplemental Table 3.

## Confocal microscopy

Standard confocal microscopy was carried out in *N. benthamiana* epidermal cells 2 to 3 days post infiltration. Cut leaf pieces were mounted on water and analyzed with a Leica DM6000B/TCS SP5 microscope (Leica Microsystems, Germany) with laser excitation settings of 488-nm for eYFP and GFP and 561-nm for RFP. Fluorescent emissions were taken at 500-550 nm for GFP and 580-620 nm for RFP. The 63x water immersion objective was used to acquire all images. Image analysis was done with the Leica LAS AF software, ImageJ and Adobe Photoshop CS5. For all Z-stacks, a distance of 1 µm was set. For induction of FLS2 internalization, flg22 peptide (EzBiolab) was gently infiltrated into leaves expressing FLS2-GFP at a final concentration of 100 µM^44^ and imaging was done 90-140 minutes post elicitation. Propidium iodide (PI, SIGMA) staining was performed as previously described^101^ with few modifications: leaf tissue was incubated in a solution of propidium iodide (200 µg/ml) for 10 minutes at room temperature, followed by three washing steps in ultra-pure water. To visualize PI-stained cells, laser excitation was set at 488-nm and fluorescence was detected between 598-650 nm.

## ACKNOWLEDGMENTS

We thank Silke Robatzek, Martina Beck and Malick Mbengue for advice to perform confocal microscopy and FLS2 internalization assays and for providing critical comments on our silencing experiments. We are grateful to Matthew Smoker for generating the AVR3a transgenic plants, Volker Lipka and Markus Albert for providing the constructs CERK1-GFP and BRI1-GFP, respectively, and Benjamin Schwessinger and Cyril Zipfel for providing the pENTR-FLS2 clone. We also thank Jacqueline Monaghan, Joe Win, Khaoula Belhaj and Yasin Dagdas for critical reading of the manuscript. This project was funded by the Gatsby Charitable Foundation, European Research Council (ERC) and Biotechnology and Biological Sciences Research Council (BBSRC).

## Supplementary Figures

**Figure S1.** All variants of AVR3a suppress flg22-mediated responses*in N.benthamiana* but show differential suppression of the marker gene *NbACRE31* after INF1 elicitation. (A, B) Total ROS production measured in relative light units (RLU) is expressed as percentage of the control treated with 100 nM flg22 over 45 minutes in transgenic plants of *N. benthamiana* (A) or *A. thaliana* (B). Plants were stably transformed with the following constructs: FLAG-AVR3a^KI^ (red), FLAG-AVR3a^EM^ (blue), FLAG-AVR3a^KI-Y147del^(grey) or vector control (ΔGFP) (green). Values are average ± SE (n = 24). Statistical significance was evaluated in comparison to the control by one-way ANOVA followed by TukeyHSD test. (C, D) Expression of the marker gene *NbACRE31* was assessed by qRT-PCR at time 0 and 180 minutes after elicitation with 100 nM flg22 (C) or 10 µg/ml INF1[Pi] (D) and normalized by NbEF1α gene expression. Results are average ± SE (n = 3 technical replicates). AVR3a variants were transiently expressed in *N. benthamiana* using the following constructs: FLAG-AVR3a^KI^ (red), FLAG-AVR3a^EM^ (blue), FLAG-AVR3a^KI-Y147del^ (grey) or vector control (ΔGFP) (green). (E, F) Western blots probed with anti-FLAG antibody after flg22 (E) or INF1 (F) treatment detected total protein expression of AVR3a variants.

**Figure S2.** AVR3a suppresses elf18-ROS production but does not affect chitin-triggered ROS accumulation. (A, B) N. benthamiana agro-infiltrated with FLAGAVR3a^KI^ (red), FLAG-AVR3aEM (blue), FLAG-AVR3a^KI-Y147del^ (grey) or vector control (ΔGFP) (green). Leaf discs were incubated in an elf18 (A) or chitin (B) containing solution and ROS was measured in relative light units (RLU) over time. Letters above the graph indicate statistical significant differences at *P* < 0.05 assessed by one-way ANOVA followed by TukeyHSD test. No statistical significance was found for group b (P = 0.068). Similar results were observed in two independent experiments.

**Figure S3.** AVR3a does not alter the subcellular localization of the surface receptors EFR or CERK1. Transient co-expression at 2.5 days post infiltration in *N.benthamiana* of EFR-YFP-HA or CERK1-GFP with FLAG-AVR3a^KI^or FLAG-AVR3a^EM^ or FLAG-AVR3a^KI-Y147del^ or vector control (ΔGFP) as indicated. Confocal microscopy shows that the plasma membrane localization of EFR-YFP or CERK1-GFP was not altered by the presence of variants of *P. infestans* effector AVR3a. Bar = 25 µm.

**Figure S4.** BRI1 constitutive internalization is not modified by AVR3a or by silencing*NbDRP2-1/2*. (A) Confocal microscopy at 2.5 days post infiltration (dpi) in *N. benthamiana* epidermal leaf cells transiently or stably expressing FLAG-AVR3a^KI^and infiltrated with BRI1-GFP. AVR3a did not alter the plasma membrane subcellular localization of BRI1 or its constitutive endosomal localization (green dots). Bar = 50 µm. Plastids auto-fluorescence (purple) is shown. (B) Quantification of the effect of AVR3a on AtBRI1-GFP endocytosis. Data was collected from three independent experiments showing similar results. The histogram depicts the number of images that showed AtBRI1-GFP endosomes as a percentage of the total number of images analyzed as indicated. Error bars are SE values. (C) BRI1-GFP was co-expressed with a hairpin-silencing construct for *NbDRP2-1/2* or the vector RNAi-GUS and confocal imaging was done at 2.5 dpi. Reduced levels of expression of *NbDRP2-1/2* did not change BRI1-GFP intracellular vesicle-like (green dots) or plasma membrane localization. Bar = 50 µm. Plastids auto-fluorescence (purple) is shown. All images are a maximum projection of 21 slices taken at 1-µm intervals. Same confocal settings were used to acquire all images. (D) Validation of *NbDRP2-1/2* silencing by RT-PCR in leaf-discs collected from the same leaves used for microscopy. (E) Quantification of the effect of RNAi-*NbDRP2-1/2* on AtBRI1-GFP endocytosis. Data was collected from 5 independent experiments showing similar results. The histogram depicts the number of images that showed AtBRI1-GFP endosomes as a percentage of the total number of images analyzed as indicated. Error bars are SE values.

**Figure S5.** *A. thaliana* dynamin GTPase DRP2A/B homologs in *N. tabacum*.ClustalW alignment shows Arabidopsis DRP2A and DRP2B dynamin GTPase proteins and homologs in *N. tabacum*. Amino acid residues are shaded dark grey if identical and a lighter shade of grey if similar. Sequences were viewed in Jalview. Full-length sequences were used for the alignment. Canonical domains previously described for large dynamin GTPase are shown.

**Figure S6.** Cladogram of dynamin related proteins (DRP). Phylogenetic tree of dynamin-related proteins (DRP) from *A. thaliana* (red), *N. benthamiana* (green), tomato (purple), potato (yellow), *N. tabacum* (dark blue), *P. trichocarpa* (pink), *V.vinifera* (black) and *M. truncatula* (light blue). The conserved GTPase-domain of DRP proteins was aligned by MUSCLE and analyzed in MEGA 6 to construct a phylogenetic tree using the maximum likelihood method. Branch length represents the estimated genetic distance. Bootstrap values for 500 replicates are shown. The sequence identifiers are from the Solgenomics, NCBI and Arabidopsis database. Green asterisks indicate the homologs of Arabidopsis DRP2A and DRP2B in *N.benthamiana* targeted by silencing in this study.

**Figure S7.** *N. tabacum*dynamin (DRP2-1) is a modular protein localized to the plasma membrane and cytosol in *N. benthamiana*. (A) Schematic representation of *N.tabacum* dynamin proteins (NtDRP2-1 and NtDRP2-2) and their domain organization: GTPase domain (G domain), Pleckstrin homology domain (PH), GTPase effector domain (GED), and a proline-rich domain (PRD). (B, C) Confocal microscopy in *N. benthamiana* epidermal cells of *Agrobacterium*-mediated expressing NtDRP2-1/2 GFP fusions. GFP-NtDRP2-1 or GFP-NtDRP2-2 primarily localized to the plasma membrane as confirmed by plasmolysis (C), far right end. (C) GFP-NtDRP2-1 and GFP-NtDRP2-2 also localized in punctuate, small vesicle-like structures. Scale bar values are shown in each picture. Representative confocal images were taken at 3 days post infiltration. Plastids auto-fluorescence (purple) is shown.

**Figure S8.** *P. capsici*effector AVR3a-4 does not associate with NtDRP2-1*in planta*. (A) *Phytophthora capsici* AVR3a-4 effector weakly co-immunoprecipitates with NtDRP2-1 *in planta*. HA-NtDRP2-1 was transiently co-expressed with FLAG-AVR3a^KI^, FLAG-PcAVR3a-4 or FLAG-RFP (control) in *N. benthamiana* and immunoprecipitated with anti-FLAG antiserum (SIGMA). Immunoprecipitates and total protein extracts were immunoblotted with the appropriate antisera. (B) Oxidative burst triggered by 100 nM flg22 in *N. benthamiana* agroinfiltrated with members of the *Avr3a* family FLAG-AVR3a^KI^, FLAG-PcAVR3a-4 or FLAG-RFP (control). ROS production was measured in relative light units (RLU) over time and depicted relative to the total ROS burst of the control. Values are average ± SE (n = 16). Statistical significance was evaluated in comparison to the control by one-way ANOVA followed by TukeyHSD test. *** P < 0.001. Experiment was repeated 3 times with similar results.

**Figure S9.** Impact of systemic silencing of*NbDRP2-1/2*in*N. benthamiana*usingVIGS. *N. benthamiana* plants were silenced using tobacco rattle virus vectors harboring a partial sequence of *NbDRP2-1* (TRV: *NbDRP2-1/2* Fragment I or TRV: *NbDRP2-1/2* Fragment II) or an empty cloning site (TRV: *GFP*). Pictures were taken 2.5 weeks after the initial infiltration with the silencing constructs.

**Figure S10.** RNAi-mediated transient silencing of*NbDRP2-1/2*in*N. benthamiana*does not compromise cell viability. (A) Schematic representation of *NbDRP2-1/2* (*Nb05397* and *Nb31648*) showing the canonical dynamin domains and the sequence region targeted by the silencing fragment. (B) Constructs carrying a hairpin plasmid (pHellsgate 8) targeting *NbDRP2-1/2* or GUS (RNAi-*NbDRP2-1/2* and RNAi-GUS, respectively) were infiltrated in *N. benthamiana* and the expression of *N.benthamiana* homologs of *AtDRP2A/B* was assessed by qRT-PCR at three days post silencing. The silencing fragment targeting part of the pleckstrin homology domain (PH, green) and the GTPase effector domain (blue) specifically knocks down the expression of *Nb05397* and *Nb31648* (*NbDRP2-1* and *NbDRP2-2*) but not the expression of *Nb11538* or *Nb09838*. Gene expression was normalized to *NbEF1α*. (C)The *NbDRP2-1/2* genes were transiently silenced in *N. benthamiana* and at three days post silencing, the epidermal cells were stained for 5 minutes with a solution of propidium iodide (PI). Auto-fluorescence of chloroplasts (purple) is shown. The absence of PI fluorescence in the cell nucleus (dashed white lines) indicates that the cells are still viable. Bar = 25 µm

**Table S1.** Plant proteins that associate with AVR3a^KI^ *in planta*.

**Table S2.** Sequence identifiers used in Figure S6.

**Table S3.** AVR3a^KI^-associated proteins identified peptides by LC-MS/MS.

